# Noda-like RNA viruses infecting *Caenorhabditis* nematodes: sympatry, diversity and reassortment

**DOI:** 10.1101/605840

**Authors:** Lise Frézal, Hyeim Jung, Stephen Tahan, David Wang, Marie-Anne Félix

## Abstract

Three RNA viruses related to nodaviruses were previously described to naturally infect the nematode *Caenorhabditis elegans* and its relative *C. briggsae*. Here we report on a collection of over 50 viral variants from wild-caught *Caenorhabditis*. We describe the discovery of a new related virus, the Mělník virus, infecting *C. briggsae*, which similarly infects intestinal cells. In France, a frequent pattern of co-infection of *C. briggsae* by the Santeuil virus and Le Blanc virus was observed at the level of an individual nematode and even a single cell. We do not find evidence of reassortment between the RNA1 and RNA2 molecules of Santeuil and Le Blanc viruses. However, by studying patterns of evolution of each virus, reassortments of RNA1 and RNA2 among variants of each virus were identified. We develop assays to test the relative potency and competitive ability of the viral variants and detect an interaction between host genotype and Santeuil virus genotype, such that the result of the competition depends on the host strain.

**Importance:** The roundworm *Caenorhabditis elegan*s is a laboratory model organism in biology. We study natural populations of this small animal and its relative *C. briggsae* and the viruses that infect them. We previously discovered three RNA viruses related to nodaviruses and here describe a fourth one, called the Melnik virus. These viruses have a genome composed of two RNA molecules. We find that two viruses may infect the same animal and the same cell. The two RNA molecules may be exchanged between variants of a given viral species. We study the diversity of each viral species and devise an assay of their competitive ability. Using this assay, we show that the outcome of the competition also depends on the host.

## Introduction

Three positive-strand RNA viruses related to nodaviruses were recently discovered to naturally infect the model organism *Caenorhabditis elegans* and its relative *C. briggsae* (1, 2). The Orsay virus (ORV) was the first natural virus found to infect *C. elegans*. Its discovery allowed to demonstrate the role of the *C. elegans* RNA interference pathway in antiviral defence (1, 3). A genome-wide association study in *C. elegans* pointed to one major locus regulating the wide range of sensitivity to the Orsay virus of *C. elegans* wild isolates: this major locus corresponds to the *drh-1 RIG-I* family gene, coding for a viral sensor triggering both small RNA and transcriptional responses (3, 4). Laboratory forward genetics also revealed a SID-3/WASP pathway necessary for viral entry (5), while 3′ terminal uridylation of viral RNAs was shown to act in viral defence (6).

In addition to the Orsay virus found in *C. elegans*, two viruses of the same family were found in *Caenorhabditis briggsae*. The Santeuil virus (SANTV) was the first to be detected (1), followed by Le Blanc virus (LEBV) (2). These viruses all infect intestinal cells (1,7) and are horizontally transmitted (1). Their genome includes two RNA segments (1, 2, 8). The RNA1 segment encodes a RNA-dependent RNA polymerase. The RNA2 segment encodes a viral capsid followed by an open reading frame (ORF) called delta. Translation of the capsid starts upstream of the first ATG and does not always end at the predicted stop codon, due to ribosomal frameshifting leading to a fusion with the delta ORF (8). The capsid structure has been solved (9). The full-length caspid-delta protein is present with low stoichiometry in Orsay virions (8), with the delta part likely protruding from the surface of the particle and possibly mediating viral entry (9). In addition, free delta appears to mediate viral exit on the apical side of the intestinal cells (10). The receptors of these viruses on the surface of the intestinal cells are still unknown. The transcriptional responses upon infection of the Orsay virus in *C. elegans* and the Santeuil virus in *C. briggsae* are in part conserved and are partially shared with the response to the infection of intestinal cells by microsporidia (11–13).

Here we aimed to collect *Caenorhabditis* viruses systematically in order to assess their diversity. *Caenorhabditis* nematodes are found in decomposing vegetal matter such as rotting fruits, stems, flowers or compost (14–16). We thereby discovered a new *C. briggsae* virus, the Mělník virus (MELV), which we show is most closely related to the Santeuil virus. We established a collection of variants of each virus, especially of the two *C. briggsae* viruses that were found most frequently, SANTV and LEBV. These viruses are often sympatric, co-existing even in the same individual worm and individual cell. Sequence analysis of the collection of viral isolates did not detect genetic exchange between these two viruses, but did detect reassortment of the two RNA molecules in SANTV. Finally, we developed a phenotypic assay that allowed us to detect interaction between the host genotype and the SANTV variant.

## Material and Methods

### Viral strain nomenclature

The names of the viruses are abbreviated as ORV for the Orsay virus, SANTV for the Santeuil virus, LEBV for the Le Blanc virus and MELV for the Mělník virus. The different viral strains were designated using the code JUv0000 where the viral strain comes from the nematode strain JU0000. In case of co-infection of the nematode strain JU0000, we used JUv0000b and JUv0000s to distinguish between the LEBV and SANTV strains, respectively. We found the viruses in different locations, including the original locations after which they are named. To avoid confusions between the viruses and the locations, we always specify ‘Santeuil virus’ when it is the virus, whereas ‘Santeuil’ alone designates the location. Finally, viral samples obtained from a population of *Caenorhabditis* coming from one rotting fruit would be coded vX0000s and/or vX0000b.

### Sampling of nematodes and associated viruses

*C. elegans* and *C. briggsae* natural populations were collected as described in (15, 17). We collected them worldwide, yet with a strong geographical bias towards France. In this area, *C. elegans* and *C. briggsae* are predominant (14, 15, 18). The presence of viruses was deduced from intestinal symptoms (1), especially when observed in the absence of other visible pathogens such as microsporidia (19) or bacteria (20), and confirmed using *in situ* hybridization (see below).

In the Orsay orchard 2013 survey, the presence of viruses was systematically monitored in *C. briggsae* using *in situ* hybridization (see below) performed directly on a subset of worms coming out of the rotten fruits. The cultures were propagated on agar plates seeded with *E. coli* OP50 and frozen in standard conditions for *C. elegans* (21).

### Detection *and de novo* sequencing of a new virus, the Mělník virus

In order to detect unknown viruses of *C. briggsae* and *C. elegans*, total RNAs were extracted from mixed-stage populations of *C. elegans* and *C. briggsae* wild isolates. These isolates were pooled, sequencing libraries constructed, sequenced on a 2×250 bp Illumina Miseq platform, and screened for viral sequences using VirusSeeker (22). Sequences related to SANTV were detected in *C. briggsae* strains JU3272 and JU3276, thereafter referred to as Mělník virus (MELV). Using RT-PCR and 3x Sanger sequencing, partial genomes of the Mělník virus RNA1 and RNA2 sequences were obtained. The genome of reference for the Mělník virus (JUv3272) is available under the accession (GenBank accession numbers MK774657, MK774659). The genome of MELV JUv3276 is available under the accession (GenBank accession numbers MK774658, MK774660).

### Detection of viruses by fluorescent *in situ* hybridization (FISH)

Each virus was detected by *in situ* hybridization using fluorescently labelled oligonucleotides specific for each virus, as described in (7). The list of probes is found in Table S1 – the use of 48/32 oligonucleotides along the viral RNA genome makes the signal brighter.

Fluorescence microscopy was performed using an upright Zeiss AxioImager M1 equipped with 10x (0.3 numerical aperture), 40x (1.3 numerical aperture), and 63x (1.25 numerical aperture) objectives. Images were acquired using a Pixis 1024B camera (Princeton instruments) and MetaVue™ imaging software. Image panels were assembled in ImageJ (23) and Inkscape (0.91 r13725, www.inkscape.org) softwares. Animals were considered infected when for an exposure below or equal to 1000 ms with the 40x objective, at least one cell was distinctly labeled at higher levels than background staining.

To screen for the natural *C. elegans* infections with ORV, we used custom Stellaris™ (Biosearch Technologies) probes labeled with Quasar® 670 dyes for the ORV RNA1 molecule and with Cal Fluor Red® 610 Dyes for the ORV RNA2 molecule.

To screen for the natural *C. briggsae* infections with SANTV and LEBV, we used custom Stellaris™ (Biosearch Technologies) probes labeled with Quasar® 670 dyes for the SANTV RNA1 and LEBV RNA2 molecules and with Cal Fluor Red® 610 Dyes for the SANTV RNA2 and LEBV RNA1 molecule. FISH was performed as described in (24), except that the hybridization solution contained 20% formamide, on each fixed sample with a 7-hour hybridization at 30°C of the Biosearch probes, targeting either the SANTV and LEBV RNA1 molecules or the SANTV and LEBV RNA2 molecules.

For viral variants’ competition experiments, we developed custom Eurofins™ 22-nt long probes specific for two divergent SANTV RNA1 variants, labeled with Texas Red for JUv1264s and with CFP-ATTO425 for JUv1993s. The probe sequences are provided in Table S1c.

### Rate of *C. briggsae* coinfection with the Santeuil and Le Blanc viruses at the fruit scale

To assess the rate of *C. briggsae* co-infection with SANTV and LEBV at the scale of the fruit, we focused on two samples from the Le Blanc location: LB14-36 (plum) & LB14-37 (pear) and one from the Orsay orchard: apple O1071. Wild hermaphrodite L4 larvae were singled less than 4 hours after the rotten vegetal matter was placed onto OP50-seeded NGM plates maintained at 23°C. After 3 days, F1 progenies were fixed with Ethanol. To monitor the presence of LEBV and SANTV viruses, FISH was performed as described in (24), except that the hybridization solution contained 20% formamide, with a 7-hour hybridization at 30°C of the custom Stellaris™ (Biosearch Technologies) probes labeled with Quasar® 670 dyes for the SANTV RNA1 molecule and with Cal Fluor Red® 610 Dyes for the LEBV RNA1 molecule (Table S1b).

### Sequencing of viral variant genomes

RNAs of a subset of infected *Caenorhabditis* wild isolates were extracted using Trizol/Chloroform. Viral genomes were first reverse-transcribed from the 3’-end, and from the middle of the segment when needed, using viral species-specific primers and the Superscript III reverse transcriptase (Invitrogen™) following the manufacturer’s instructions. The reverse transcription was followed by PCR amplifications using the Q5® high-fidelity DNA polymerase (New England Biolabs) with primers amplifying two overlapping fragments per RNA segment. The PCR products were Sanger sequenced. To minimize sequence errors due to the PCR, we performed two independent reverse transcription-PCR but did not find any mismatches between replicates and were able to validate heterozygous SNPs. All the primers used in this study are listed in Table S1a. Sequences are available under the accessions xxxx.

### Analyses of viral genetic diversity

#### Alignments and phylogenetic analyses

Nucleotide sequences were aligned using MUSCLE (25) with default parameters implemented in MEGA7 (26). When necessary, heterozygous sequences were phased using fastphase implemented in DNAsP v5 (27). The sequence relationships were inferred in MEGA7 (26) using the Maximum Likelihood method based on the Jones-Taylor-Thornton (JTT) matrix-based model (28) and tested using 10,000 bootstraps. Initial tree(s) for the heuristic search were obtained automatically by applying Neighbor-Join algorithms to a matrix of pairwise distances estimated using a JTT model, and then selecting the topology with superior log likelihood value. Timetrees of SANTV and MELV phylogenetic relationships were inferred using the Reltime method (29) and estimates of branch lengths inferred using the Neighbor-Joining method (30).

#### Detection of reassortments

To detect events of RNA molecules’ reassortment between variants of the same viral species, we concatenated, for each variant, their RNA1 and RNA2 nucleotide sequences. We detected putative reassortment and recombination events using the RDP, GENECONV, Bootscan, Maxchi, Chimaera, SiSscan, 3Seq methods implemented in RDP4 (31) software. For all these methods, the following parameters were used: neighbor-joining tree built using 1,000 bootstraps; recombination events were considered significant for a p-value lower than 0.001.

#### Analysis of polymorphisms along the ORFs within the set of LEBV and SANTV variants

Mean evolutionary diversities for the entire populations and mean interpopulational evolutionary diversities were estimated using MEGA7 (26), following Nei and Kumar calculations (32). We calculated the number of base substitutions per site and the number of amino acid/or nucleotide substitutions per site (i) from mean diversity calculations for the entire population, and when specified from mean interpopulational diversity. Standard error estimates were obtained by a bootstrap procedure (1,000 replicates). Analyses were conducted using Maximum Composite Likelihood model (33) for nucleotides and using the JTT matrix-based model (28) for Amino acids. All positions with less than 90% site coverage were eliminated, i.e. fewer than 10% alignment gaps, missing data, and ambiguous bases were allowed at any position.

The total number of mutations (S) and the ratios of non-synonymous to synonymous substitutions (dN/dS), where dS is the number of synonymous substitutions per site (s/S) and dN is the number of nonsynonymous substitutions per site (n/N), were calculated for the RNA1 (RdRP) and RNA2 (Caspid-delta fusion protein) molecules of SANTV and LEBV in a sliding-window, with a window of 20 bp and a step of 1 bp. In order to describe the distribution of the total, synonymous and nonsynonymous polymorphisms we used the maximum likelihood analysis of natural selection codon-by-codon using the HyPhy software package (34) implemented in MEGA5 (35). For each codon, estimates of the numbers of inferred synonymous (s) and nonsynonymous (n) substitutions, the numbers of sites that are estimated to be synonymous (S) and nonsynymous (N) were produced using the joint Maximum Likelihood reconstructions of ancestral states under a Muse-Gaut model (36) of codon substitution and Felsenstein 1981 model (37), of nucleotide substitution. For estimating ML values, a tree topology was automatically computed. To detect codons that have undergone positive selection, we used the [dN – dS] statistical test implemented in the HyPhy software package (34). The dN/dS ratio values were then calculated for a sliding-window with a window of 20 bp and a step of 1 bp.

The codon-based tests of positive and purifying selection averaging over all sequence pairs were conducted in MEGA7 (26), providing the probabilities of rejecting the null hypothesis of strict-neutrality (dN = dS) in favour of the alternative hypothesis (dN > dS) =1.00 or (dN < dS) =1.00. The variance of the difference was computed using the bootstrap method (10,000 replicates). Analyses were conducted using the Nei-Gojobori method (38). All ambiguous positions were removed for each sequence pair.

### Viral preparations

Naturally infected *C. briggsae* (*Cbr*) wild isolates were thawed and propagated for 7 days onto OP50 seeded NGM plates at 23°C. Viral preparations were made from these cultures as in (3) for the following viruses: SANTV (JUv1264, JUv1993, JUv1551), LEBV (JUv1498), ORV (JUv1580, JUv2572) and MELV (JUv3272).

### Comparison between ORV, SANTV and LEBV viral potencies

First, we established *C. elegans* (*Cel*) infections with two ORV variants. Five L4 larvae were infected with 50uL of viral preparation. Infected *Cel* populations were propagated onto OP50 seeded NGM plates at 20°C for 2-3 host generations so that the final titer is function of the host-pathogen interaction rather than of the initial titer (1, 3). The original host strain *Cel* JU1580, and three other *Cel* strains N2, JU2572 and MY10 were infected with the ORV JUv2572 strain from Ivry and with the original ORV JUv1580 isolate from the Orsay orchard (Figure S6). F2/F3 populations were harvested and fixed for *in situ* hybridization using custom Stellaris™ probes targeting ORV RNA2 molecules (Table S1b). The number of infected cells was counted manually.

Second, we performed pairwise competition assays between SANTV JUv1993 and two other SANTV variants, JUv1264 and JUv1551. We independently infected *Cbr* JU1264 animals with the tested viral preparations (i.e. JUv1993, JUv1264 and JUv1551) in three replicates. We checked that after 7 days, more than 80% of the nematodes were infected. We then mixed 20 (10 + 10) sick adult animals (visual symptoms) from the originally infected populations with either of two viruses and let them reproduce for 3 days at 23°C. We passed at each generation (every 3 days) 20 L4 larvae to a new plate and fixed each generation for *in situ* hybridization using 22nt-specific probes specified in Table S1c. The number of (co-) infected cells was counted manually.

From the above experiment, we also prepared a virus filtrate in the first generation of co-infection, and re-infected several *Cbr* wild isolates (JU1264, JU1498, JU1993, JU2160). We then passed at each generation (every 3 days) 20 L4 larvae to a new plate and fixed each generation for *in situ* hybridization using 22nt-specific probes specified in Table S1c.

### Tissue tropism of the Mělník virus in *C. briggsae*

Cultures of naturally infected JU3272 and JU3276 *Cbr* isolates were propagated for 5 days onto *E. coli* OP50 seeded NGM plates at 23°C before the fixation of mixed-stage populations. We also infected five L4 larvae of the *Cbr* strain JU1498 with 50 µL of the JUv3272 viral preparation. Infected cultures were propagated for 5 days onto OP50-seeded NGM plates at 23°C before the fixation of mixed-stage populations.

The tissue tropism of MELV was characterized using fluorescent *in situ* hybridization. We developed a custom Eurofins™ 21-nt long probe labeled with Texas Red dye for the MELV RNA1 molecule (Table S1c). FISH was performed on fixed animals as described by (24), except that the hybridization solution contained 20% formamide.

## Results

### The Mělník virus: a new noda-like virus infecting *Caenorhabditis briggsae*

We sampled *C. elegans* and *C. briggsae* and searched for possible viral infections using *in situ* hybridization with probes corresponding to known viruses and whole-RNA sequencing of infected cultures (see Methods). A novel *C. briggsae* virus, Mělník virus (MELV), was found in Mělník and Prag (Czech Republic). This virus infects intestinal cells (Figure S1) and is not transmitted vertically, as shown by the loss of infection after submitting the culture to a bleach treatment. MELV is related to the three previously identified noda-like viruses. Partial genome sequences were obtained of 2342 nt for the RNA1 and 2257 nt for RNA2 which encompass a portion of the CDS for the RdRP and likely the entire Capsid-delta proteins, respectively. We aligned the amino-acid sequences of the RNA-dependent RNA polymerase (RdRP) and capsid-delta proteins of the four viruses (Figures S2 and S3; Figure 1B for their pairwise divergence). Both the RdRP and capsid sequences of this new virus are most related to those of the Santeuil virus (Figure 1B-D).

**Figure 1.**
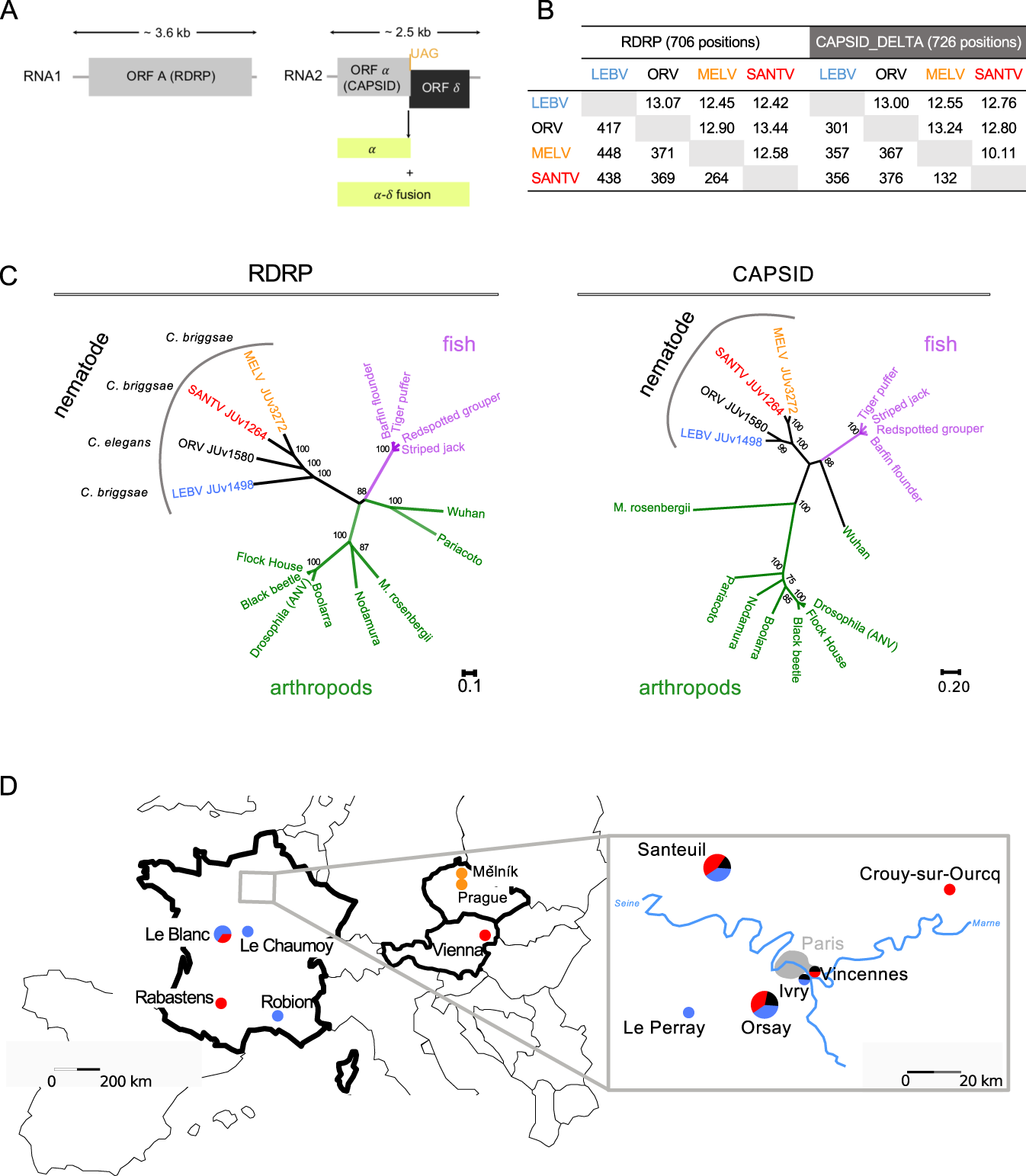
Phylogenetic and geographical distribution of the four sampled *Caenorhabditis* viruses, including the new Mělník virus. **(A)** Genome structure of the four noda-like *Caenorhabditis* viruses. **(B)** Pairwise number of amino acid differences between noda-like *Caenorhabditis* viruses (below the diagonal) and standard error estimates (above the diagonal). All positions containing gaps and missing data were eliminated. **(C)** Phylogenetic relationships between the RdRP and capsid amino-acid sequences of classic nodaviruses and the four noda-like *Caenorhabditis* viruses. The Delta ORF is not found in the classical nodaviruses. **(D)** Geographical distribution of the sampled viruses. The viruses are colour-coded: Orsay virus (ORV) in black, Santeuil virus (SANTV) in red, Le Blanc virus (LEBV) in blue and Mělník (MELV) in orange. For a detailed list of strains, see Table S2.

### Host specificity and geographical distribution of the four viruses

In our surveys in France and the occasional sampling elsewhere in Europe, *C. elegans* was only found infected by ORV. *C. briggsae* was found infected by SANTV, LEBV or MELV, but never by ORV. We did not find intestinal infections by viruses in other *Caenorhabditis* species. This pattern of host specificity in natural populations matches the pattern of infection in the laboratory ((1); and our results). For example, SANTV, LEBV and MELV all infect *C. briggsae* JU1498 but do not infect the *C. elegans* ORV-sensitive strain JU1580. Conversely, the ORV variants JUv2572 and JUv1580 infect various *C. elegans* isolates (see below) but do not infect *C. briggsae* JU1264.

Although *C. elegans* and *C. briggsae* were found at similar frequencies in France, viral infections of *C. briggsae* were easier to find than *C. elegans* infections (Fig. 1). Finding infected *C. elegans* was indeed rare, even in the highly resampled Orsay orchard. We only found it again in the Orsay orchard in 2014 and once in two other locations, Ivry and Santeuil (Figure 1; Table S2).

*C. briggsae* is commonly found worldwide (16), yet we rarely found signs of infection outside France, and never outside Europe despite *C. briggsae* cosmopolitan occurrence (14). Our virus set is thus highly geographically biased towards France. In France, we found SANTV and LEBV at similar frequencies in *C. briggsae*. The two viruses were found co-existing in the three most sampled locations: we found LEBV in the Santeuil wood, SANTV in Le Blanc and both in the Orsay apple orchard (Figure 1D). Over the 2008-2014 period, we established from the Orsay orchard 20 *C. briggsae* isolates (strains or F1 progenies; see Methods and Figure 2A) with SANTV only, 13 with LEBV only and 10 with both. Including the other locations, we established 26 cultures with the SANTV only, 27 with LEBV only and 12 with both (Figure 1, Table S2).

**Figure 2.**
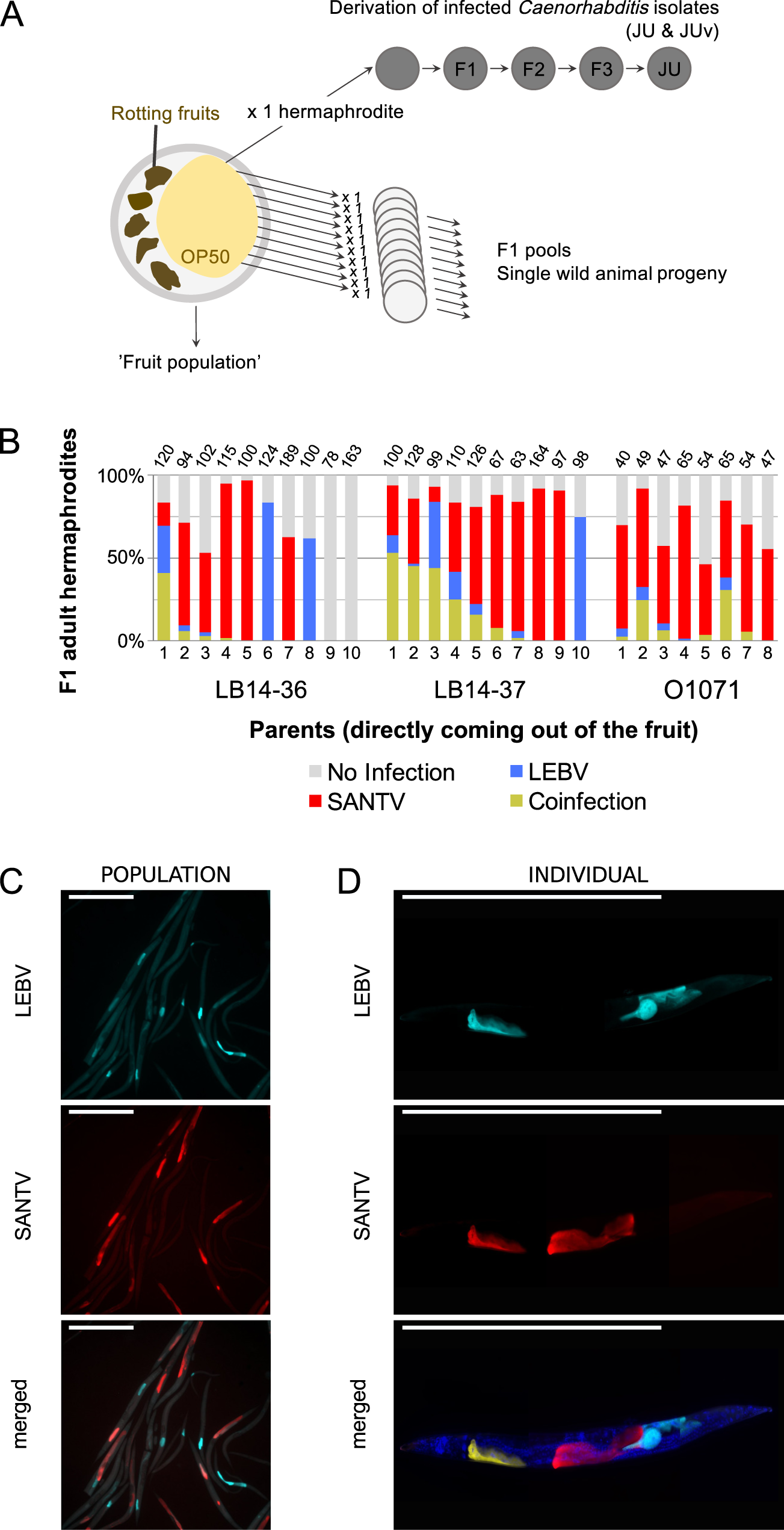
Co-infection of *C. briggsae* by Santeuil and Le Blanc viruses at the level of population, individual animals and single cells. **(A)** Design of the assay. Rotten vegetal matter was collected from the wild, brought back to the laboratory and plated on *E. coli* OP50 seeded NGM plate. Single hermaphrodites of *C. briggsae* were isolated onto a new plate. *C. briggsae* wild isolates (JU) naturally infected with their noda-like viruses (JUv) were derived from the ‘fruit population’. In the survey for co-infection (panel B), subsets of the *C. briggsae* wild populations coming out of the fruit (’fruit population’) were fixed in ethanol and the presence of viruses was systematically monitored using *in situ* hybridization. **(B)** Natural co-infection in the progeny of wild animals (numbered on the *x* axis) isolated from rotten fruits. Three samples where the SANTV and LEBV were present were chosen to assess the co-existence of the two viruses at the level of individual worms: a plum (LB14-36) and a pear (LB14-37) from the Le Blanc location and an apple (O1071) from the Orsay orchard. The F1 progeny (‘F1 pool’) of 8-10 parents isolated from the parental populations were fixed in ethanol when they reached the adult stage. Many of the isolated adults harbored both viruses as assayed by their presence in their progeny and some of their progeny also detectably harbored both viruses. Note that in all three cases the Santeuil virus appears predominant. The number of scored animals in each F1 pool is indicated on top of the bar.Co-infection at the level of the single *C. briggsae* LB14-37.1 F1 population. Fluorescence *in situ* hybridization staining of SANTV and LEBV-infected animals using SANTV RNA1 (Quasar® 670) and LEBV RNA1 (Cal Fluor Red® 610) Biosearch® probes. Nuclei were counterstained with DAPI (merge panels). Red: SANTV RNA1 probe; Cyan: LEBV RNA1 probe; Scale bars represent 500 μm. **(D)** Co-infections were recorded at the level of a single individual and a single cell both in naturally and in artificially infected *C. briggsae* animals. Here, the illustration of a *C. briggsae* adult hermaphrodite (strain JU1264) artificially coinfected with JUv1264 and JUv1498 in the laboratory. Fluorescence *in situ* hybridization staining of SANTV and LEBV-infected using SANTV RNA1 (Quasar® 670, 40x, 200ms) and LEBV RNA1 (Cal Fluor Red® 610, 40x, 200ms) Biosearch® probes (see Table S1b). Nuclei were counterstained with DAPI (merge panels). Red: SANTV RNA1 probe; Cyan: LEBV RNA1 probe; Yellow: for co-infected cells. Scale bars represent 500 μm.

### Co-occurrence of SANTV and LEBV in *C. briggsae*

We focused on sampling *Caenorhabditis* viruses in the Orsay orchard, where nematodes from a large set of 226 apples were systematically surveyed from 2010 to 2014 by *in situ* hybridization for the presence of the SANTV and LEBV viruses. In this set, 91/221 apples contained *C. briggsae*, and 43 of those were infected by a virus; out of the infected isolates, 20/43 were infected by SANTV only, 13/43 with the LEBV only, and 10/43 were co-infected by both viruses. Co-infection of *C. briggsae* ‘fruit populations’ was also found in Le Blanc in 2014.

In order to assess the rate of *C. briggsae* co-infection at the scale of individual wild-caught animals, we then focused on three naturally co-infected populations (a pool of *Cbr* animals coming out of the rotting fruits LB14-37, LB14-36 and O1071; Figure 2). The co-infection rates in the progeny (F1) greatly differed among F0 individuals, ranging from no detectable infection to 50% of co-infection (Figure 2B; example of co-infection signal in a F1 population shown in Figure 2C). In the highly-infected population, LB14-37, 7 animals out of 10 had more than 10% of their progeny co-infected. In the moderately-infected populations, LB14-36 and O1071, only 1/10 and 2/8 animals had more than 10% of their progeny co-infected, suggesting that the co-infection of a ‘fruit population’ can be lost relatively quickly when establishing an infected *Caenorhabditis* isolate (JU and its virus-es-JUv) as in Figure 2A.

We next asked whether the two viruses could be found in the same individual nematode and the same intestinal cell. Our results show that SANTV and LEBV could be found infecting the same cell in naturally infected *C. briggsae* animals (Figures 2B,C) as well as in *C. briggsae* isolates simultaneously infected by both viruses under laboratory conditions (Figure 2D).

### Genetic diversity, phylogeny and reassortments

We obtained by Sanger sequencing the near-entire RNA1 and RNA2 segments for a large set of the collected viruses (listed in Table S1a). We did not find any evidence of genetic exchange between the different *Caenorhabditis* noda-like viruses. Indeed, the SANTV viruses always contained both RNA1 and RNA2 molecules that were closely related to our original SANTV virus strain, JUv1264 (1) and similarly for LEBV viruses (2).

Within each viral species, we found extensive genetic diversity among the strains. The highest genetic diversity was found for the SANTV RNA1 molecule with a total of 21% nucleotides (11.9% amino-acids) showing a polymorphism in the whole sequenced set. Mean pairwise diversity for the entire sample was 0.086±0.019 (standard error, SE) substitutions per site at the nucleotide level and of 0.035±0.005 at the amino-acid level. The pattern of SANTV RNA1 diversity includes a deep split between two lineages, which we call A and B (Figure 3, Figure S6). The estimate of evolutionary divergence over sequence pairs between RNA1 of lineages A and B was 0.099±0.037. The RNA1 lineages A and B had similar mean diversity 0.0024±0.002 and 0.0024±0.007 substitutions per site at the nucleotide level and of 0.012±0.002 and 0.014±0.002 at the amino-acid level, respectively. Lineage A was found in Orsay in 2008, 2009, 2010, 2012 and 2013 and in several other locations, while lineage B was found in Orsay in 2010 and 2013 and Vienna in 2017.

**Figure 3.**
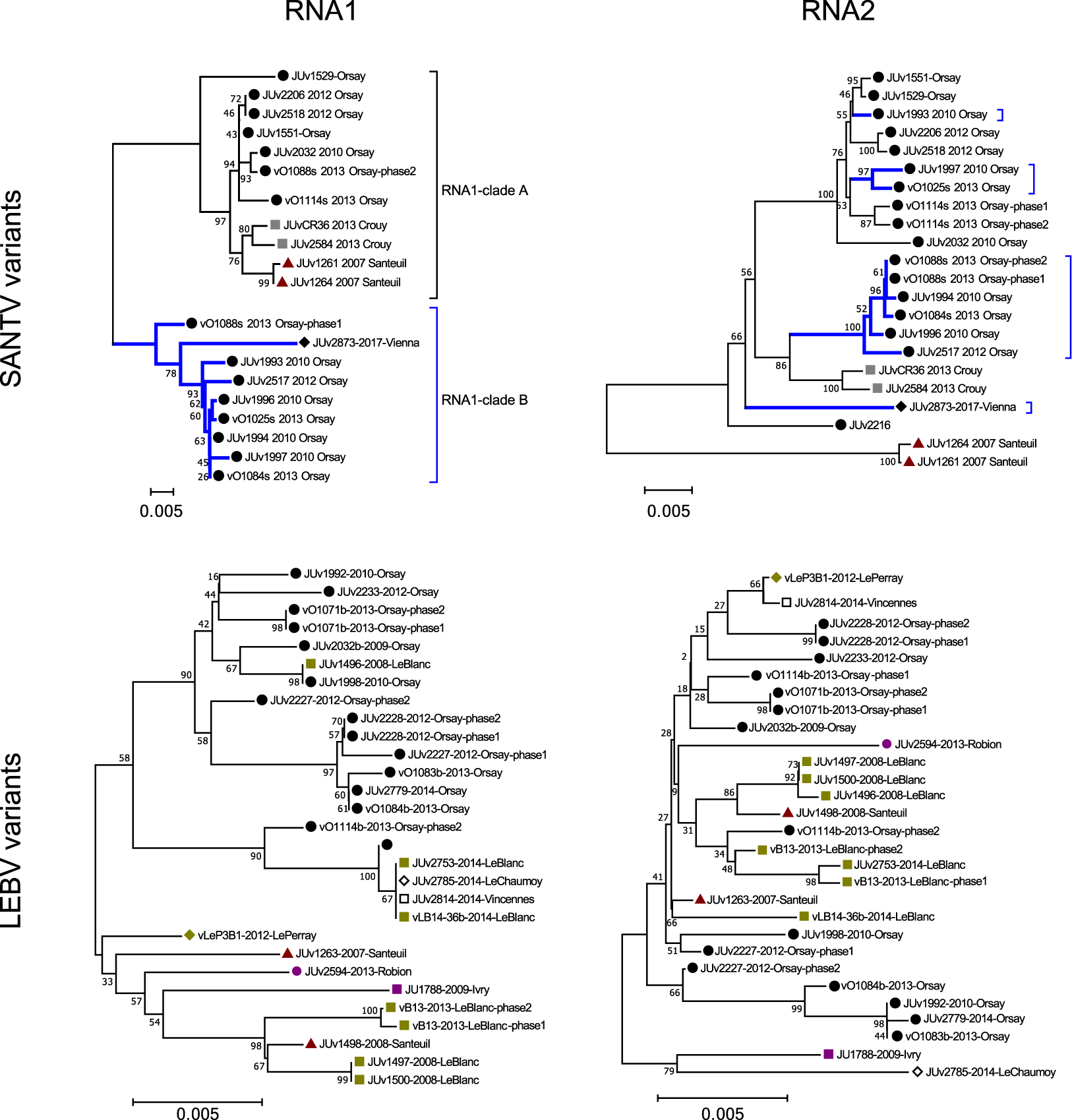
Phylogenetic relationships of the SANTV and LEBV variants. The top panels show the RNA1 and RNA2 trees for SANTV, the bottom panels for LEBV. The phylogenetic relationships were inferred using the Neighbor-Joining method (see Methods). All positions containing gaps and missing data were eliminated. The percentage of replicate trees in which the associated taxa clustered together in the bootstrap test (10,000 replicates) are shown next to the branches. The evolutionary distances were computed using the Maximum Composite Likelihood method. The tree is drawn to scale with the number of base substitutions per site. The locations are coded by a colored shape next to the name of the variant. For SANTV, the branches leading to variants in the clade B of RNA1 are labelled in blue. These variants do not cluster in the RNA2 tree, indicating reassortment between RNA1 and RNA2. All positions with less than 90% site coverage were eliminated. The total of positions used to build trees were of 2877 nt for SANTV RNA1, 2251 nt for SANTV RNA2, 2976 nt for LEBV RNA1 and 2523 nt for LEBV RNA2.

The SANTV RNA2 molecule displayed a lower level of variation than RNA1: mean diversity for the entire sample was 0.027±0.002 (SE) substitutions per site at the nucleotide level and 0.011±0.002 at the amino-acid level. Interestingly, the phylogeny of RNA2 was not congruent with that of RNA1, suggesting possible reassortments (Figure 3, variants indicated by the blue lines pointing to them). Using RDP4 (31), we indeed detected five reassortment events between the RNA1 and RNA2 molecules (Figure 4; Table S3). No significant intramolecular recombination was detected in our panel of viral variants. We did find populations from a single fruit that were infected by two variants (Figure 3). Thus, it is likely that SANTV virus variants reassort while co-infecting the same individual nematode.

**Figure 4.**
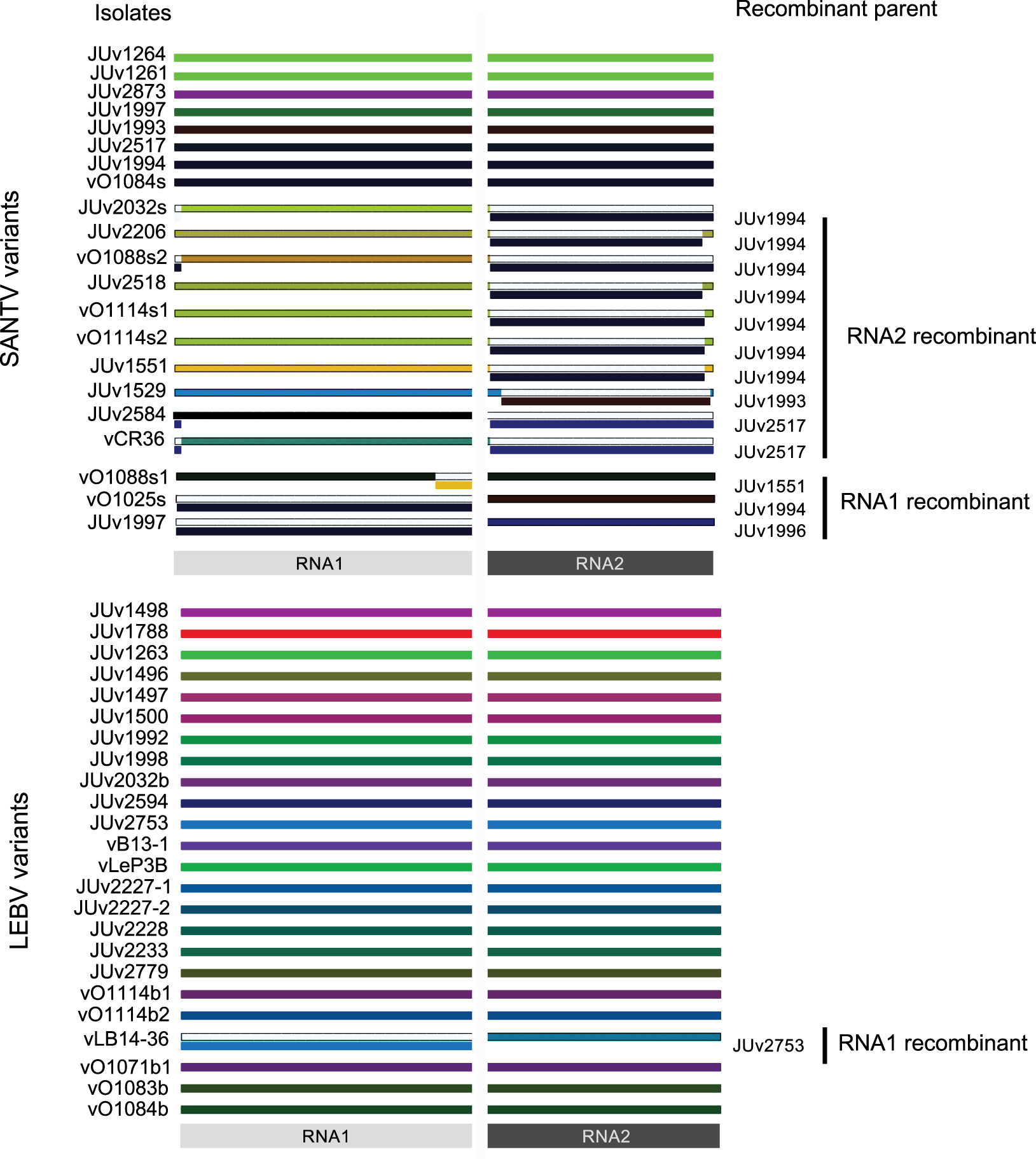
Reassortment of RNA1 and RNA2 molecules. Output of RDP4 analysis. Top panel: SANTV wild isolates. Bottom panel: LEBV wild isolates. Names of the wild viral isolates are specified on the left. Each color indicates a haplotype. When reassortment is detected, the name of the putative haplotype donor (or recombinant parent) is specified on the right and a line with the parent haplotype’s color is shown under the wild isolate haplotype white line. The *p*-value corresponding to the rejection of the hypothesis of absence of reassortment or recombination are given in the Table S3.

For LEBV, despite a similar sampling structure and geographical diversity, we found a much lower level of genetic diversity than for the Santeuil virus on both RNA molecules. The RNA1 molecule displays 0.013±0.001 substitutions per site at the nucleotide level and 0.007±0.001 at the amino-acid level, while the RNA2 harbors 0.011±0.001 substitutions per site at the nucleotide level and 0.007±0.001 at the amino-acid level. The topology of the phylogenetic relationships between LEBV variants for RNA1 and RNA2 was poorly supported, possibly because of the lower number of informative sites (Figure 3). Using RDP4 (31), we detected one event of reassortment between RNA1 and RNA2 molecules (Figure 4; Table S3).

The pattern in the number of polymorphic sites (S) and the ratio of non-synonymous to synonymous substitutions (dN/dS) along the RNA1 and RNA2 molecules are shown for the LEBV and SANTV variants in Figure 5A. Perhaps most striking is the low level of synonymous polymorphism compared to non-synonymous polymorphisms around codon 725 of SANTV RNA1 and 70 of LEBV RNA1. The [dN – dS] statistical test implemented in the HyPhy package (34) did not however detect any positive selection in our data set. The Z-test (codon-based tests of positive and purifying selection for analysis averaging over all sequence pairs) rejected the null hypothesis of strict-neutrality in favor of purifying selection for the RNA1 and RNA2 of both LEBV and SANTV.

**Figure 5.**
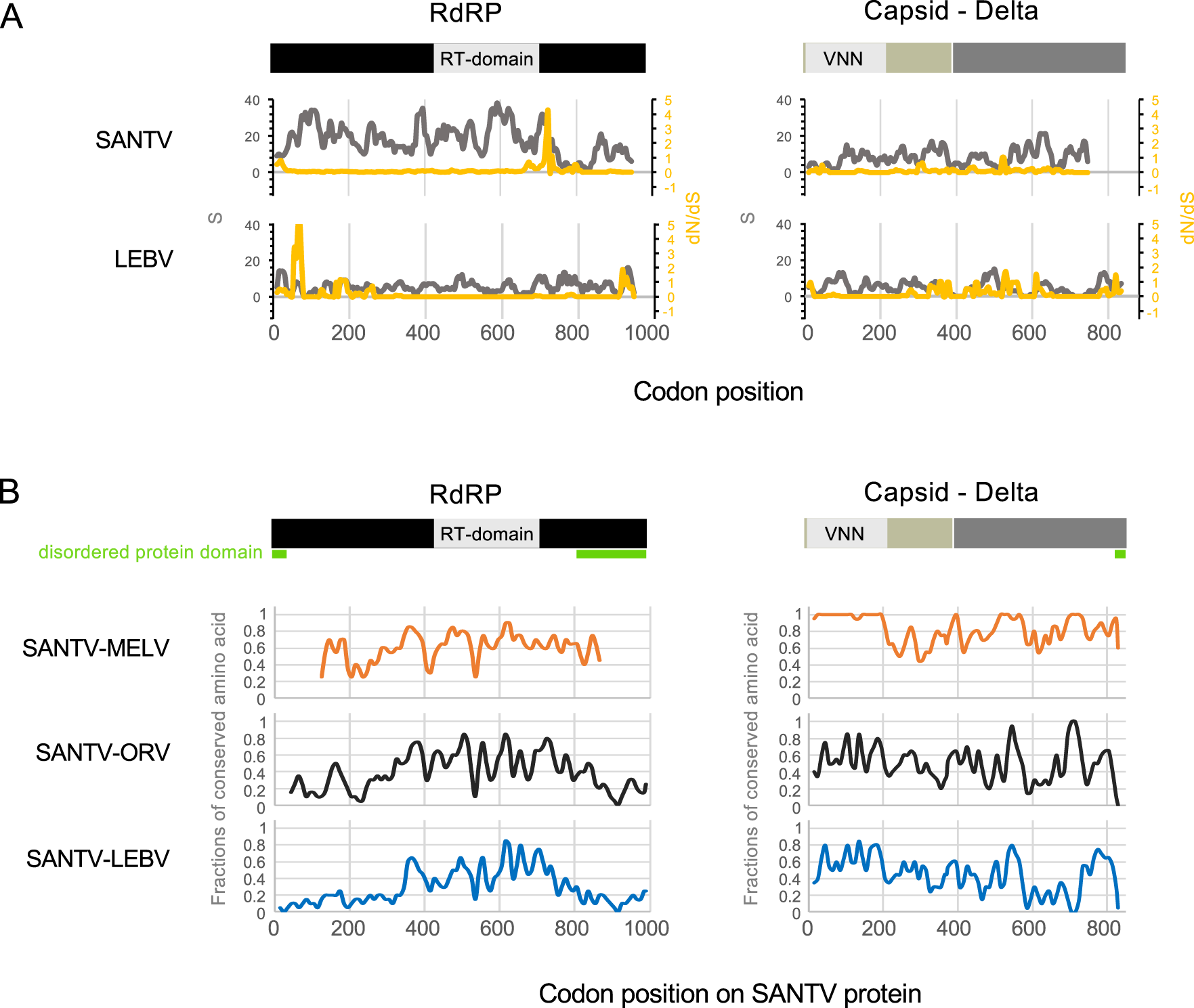
Pattern of molecular diversity along the open reading frames. **(A)** The total number of mutations (S) are indicated in grey and the ratios of non-synonymous to synonymous substitutions (dN/dS) in yellow, where dS is the number of synonymous substitutions per site (s/S) and dN is the number of nonsynonymous substitutions per site (n/N). Values were calculated for the RNA1 (RdRP) and RNA2 (Caspid-delta fusion protein) molecules of SANTV and LEBV in a sliding-window, with a window of 20 codons and a step of 5 codons. (**B)** The fractions of amino acid conserved between SANTV and the other noda-like viruses (MELV, ORV and LEBV) along the RNA1 and RNA2 molecules were estimated in a sliding window (20AA:10AA). RNA-dependent RNA polymerase (RdRP) is in black, including the RT (reverse transcriptase) domain. Capsid is in beige, including the VNN (viral nervous necrosis) domain that belongs to the viral-coat superfamily (S domain), and Delta is in dark grey. Disordered protein domains as predicted by the PrDOS protein disorder prediction server (http://prdos.hgc.jp) are underlined in green.

Finally, the pattern of pairwise amino-acid conservation between SANTV, MELV, ORV and LEBV along the RNA1 and RNA2 molecules are shown in Figure 5B. No amino acid region is specifically conserved between the three *C. briggsae* viruses (SANTV, MELV, LEBV) that is not also shared with the *C. elegans* virus (ORV). Remarkably, the N-terminal capsid domain is highly conserved between the two closely related viruses MELV and SANTV.

### Phenotypic assays of potency of the viral variants

We observed that some host-virus combinations caused more damage to the host than others, as revealed by intestinal symptoms and slow growth (data not shown). We previously reported that *C. elegans* wild isolates display a wide range of sensitivity to a given viral strain (3) and will report elsewhere on the diversity found in *C. briggsae* (see also (1)). Here we aimed to compare the virus variants and test the effect of genetic diversity within viral species on their potency in a given host. The RT-PCR titer of our filtered viral preparations may not reflect the amount of infectious viral particles. To circumvent this titration problem, we used several protocols that aimed at measuring viral production by a given host independently of the original viral titer.

First, in the simplest protocol, we established infections and monitored the amount of virus produced after 2-3 host generations so that the final titer is function of the host-pathogen interaction rather than of the initial titer (1, 3). Using this protocol, the ORV JUv2572 strain from Ivry appeared more potent than the original ORV JUv1580 isolate when tested on the original host strain JU1580 as well as on strain MY10, which is resistant to infection by JUv1580 (3) (Figure S6A-B). Moreover, JUv2572 tends to infect its host intestine more anteriorly than JUv1580, and this was independent of the *C. elegans* isolate infected (Figure S6C-D). This new ORV isolate may constitute a useful resource for *C. elegans* viral infections.

Second, in order to best account for competitive fitness of the viruses infecting *C. briggsae*, we directly tested the competitive ability of two SANTV strains in the same host, using a two-step infection protocol (see Methods and Fig. 6A). We pre-infected *C. briggsae* JU1264 with each of three tested viral preparations in three replicates in conditions such as after 7 days (2-3 host generations), more than 80% of the nematodes were infected. Then we mixed 10 sick adult animals of the co-infected populations and let them reproduce for 3 days. We then passed every 3 days 20 L4 larvae from the population for three generations. In each generation, ∼ 100 animals were then fixed for *in situ* hybridization.

**Figure 6.**
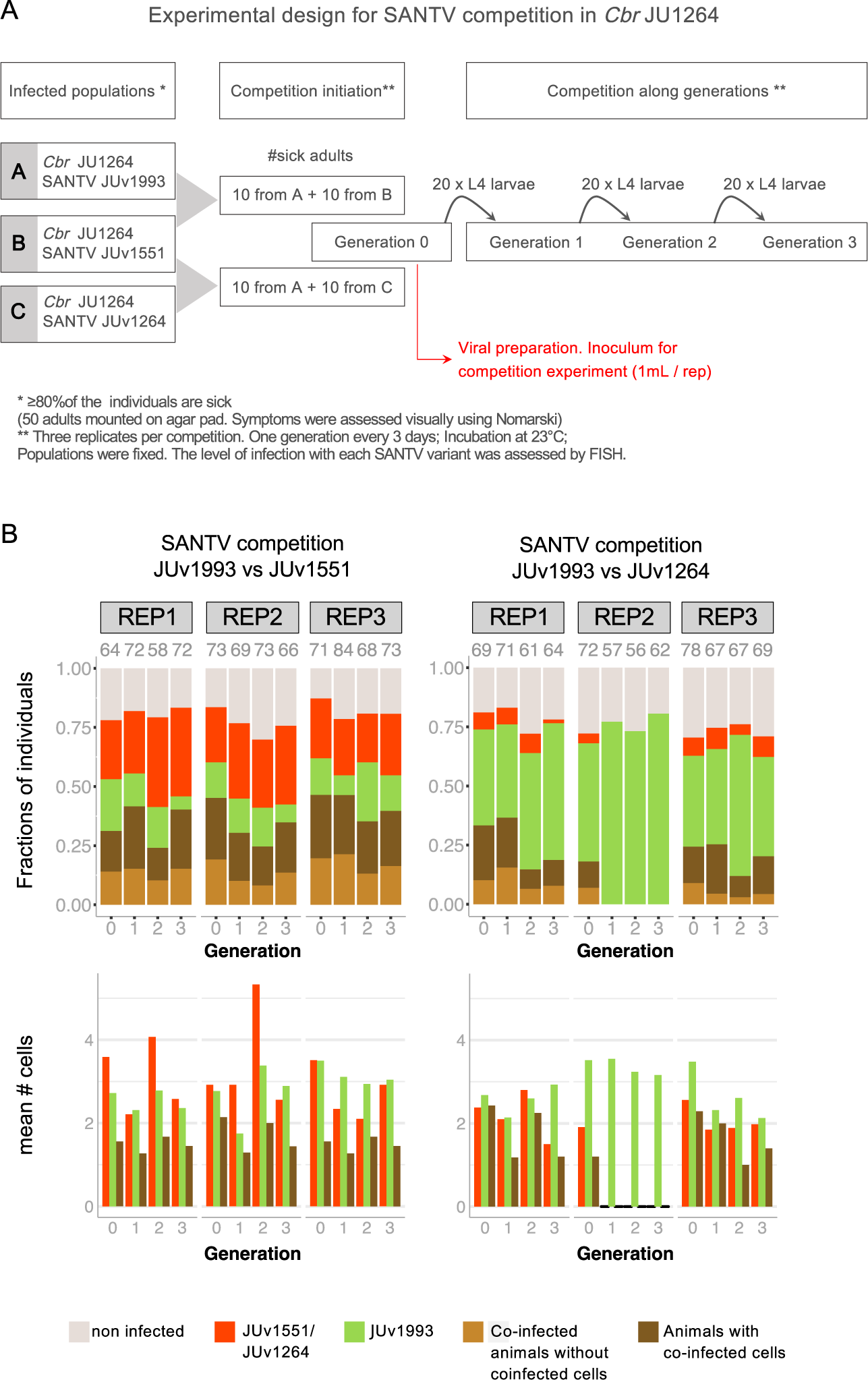
Pairwise infections of *C. briggsae* JU1264 by three SANTV variants. **(A)** Design of the assay for the pairwise competition between the JUv1993 SANTV variant and each of the two SANTV variants JUv1264 (variant of reference) and JUv1551. JUv1993 and JUv1551 were chosen as they possess highly divergent RNA1 and similar RNA2 molecules (see Figure 3). The result is shown in panel B. The viral preparation (in red) is then used for the experiment in Figure 7. **(B)** Result of the above experiment in *C. briggsae* JU1264, expressed as a fraction of infected individuals (top graphs) and mean number of infected cells in the infected animals (bottom graphs), across four generations. The Santeuil virus variants JUv1551 and JUv1993 were maintained at similar frequencies (left), whereas JUv1264 tends to lose to JUv1993 (right). ‘1’, ‘2’, ‘3’ designate the generation of passage of the population. ‘REP’: replicate. (Co-) infection levels were scored using FISH staining of SANTV infected cells/animals using single oligonucleotide probes targeting JUv1264/JUv1551 RNA1 (ATTO425) and JUv1993 RNA1 (Texas Red) (see Table S1c). The total number of animals scored per replicate are given above the histograms.

To follow two SANTV variants, we developed probes specific for two divergent SANTV RNA1 from the two clades in Figure 3 (Table S2; we do not know whether RNA1-RNA2 reassortments may have occurred during the experiment). With this assay, we observed reproducible differences between SANTV genotypes. JUv1551 and JUv1993 RNA1 were both maintained quite stably over the four generations and both infected similar proportions of infected animals and number of cells. In the JUv1993 versus JUv1264 competition, more animals were infected by JUv1993 RNA1 than by JUv1264 RNA1; the latter even disappeared by the second generation in one replicate (Fig. 6B). We conclude that JUv1993 outcompetes JUv1264 in the *C. briggsae* JU1264 host.

Third, we sought to further test the competitive ability of these viruses in several *C. briggsae* host strains, by starting a new infection from the mixed viral preparation in the first generation of co-infection from the previous experiment (in red in Fig. 6A). We infected in parallel *C. briggsae* strains JU1264 (originally infected by SANTV), JU1498 (by LEBV), JU1993 (by SANTV) and JU2160 (tropical isolate from Zanzibar). Consistent with the above result, the Santeuil virus genotypes JUv1551 and JUv1993 were co-maintained in *C. briggsae* JU1264 in similar proportions of infected animals across the three generations, while more animals were infected by JUv1993 than by JUv1264 (Figure 7). The same result applied when the JU1993 host was infected. Interestingly, the viral genotype JUv1993 appeared to outcompete JUv1264 better in the host strain JU1498 than in JU1264 (glm using a logistic regression on the number of cells respectively infected with JUv1993 and JUv1264, and a quasibinomial model with logit link, p=0.006) and even better in the host strain JU2160, where the JUv1264 variant was lost in all three replicates by the third generation (glm using a logistic regression on the number of cells respectively infected with JUv1993 and JUv1264, and a quasibinomial model with logit link, p=0.024).

**Figure 7.**
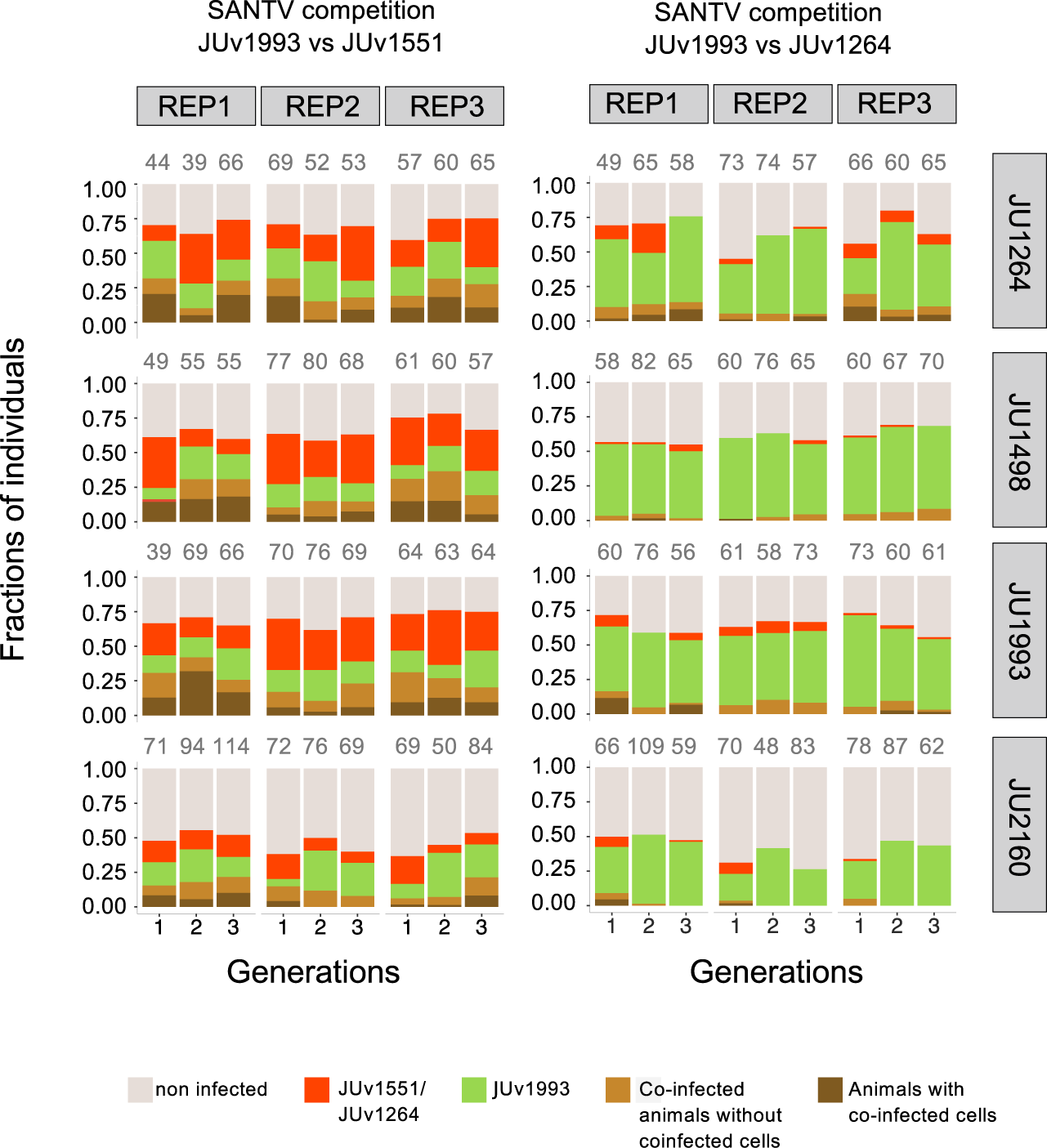
The outcome of competition between two SANTV variants depends on both the Santeuil virus genotypes and the host genotype. The tested SANTV genotypes are indicated above the graph and the *C. briggsae* host genotype on the right. The different *C. briggsae* strains were infected from viral preparations prepared as in Fig. 6A. ‘1’, ‘2’, ‘3’ designate the generation of passage of the population, generation 1 being the progeny of the originally infected animals. All experiments were performed in parallel. (Co-) infection levels were scored using FISH staining of SANTV infected cells/animals using single oligonucleotide probes targeting JUv1264/JUv1551 RNA1 (ATTO425) and JUv1993 RNA1 (Texas Red) (see Table S1c). The total number of animals scored per replicate are given above the histograms

In summary, these experiments indicate i. a different competitive ability of different viral strains in a given host and ii. an interaction between host genotype and Santeuil virus genotype, such that the result of the competition depends on the host strain.

## Discussion

To date, four *Caenorhabditis* noda-like viruses have been discovered. The Santeuil, Orsay, Le Blanc viruses and the new Mělník virus share the same tissue tropism towards intestinal cells, and are horizontally transmitted. The capsid-delta protein is more conserved among them than the RdRP protein. The newly discovered Mělník virus is closely related to the Santeuil virus. The MELV RNA2 molecule is particularly close to the SANTV RNA2 molecule, especially in the 207 first amino acids of the capsid protein, suggesting that the RNA2 molecule (capsid-delta protein) is evolutionarily more constrained than the RNA1 molecule (RdRP). While the ORV virus infects *C. elegans* and the other three viruses infect *C. briggsae*, the ORV and LEBV capsid-delta proteins share more similarities between each other than with SANTV and MELV. The amino-acid conservation and phylogenetic positions of the four viruses suggest that ORV has derived from LEBV virus while specializing on the host *C. elegans*. Further comparison between LEBV and ORV including reconstitution of infection through transgenes as in (8) could therefore provide a better understanding of the host switch from *C. briggsae* to *C. elegans*.

Although *C. elegans* and *C. briggsae* are found at similar frequencies in France (17,18) we mostly recorded viral infections of *C. briggsae*. The number of viral infections recorded for *C. elegans* was low even if, under laboratory conditions, many European *C. elegans* wild isolates are sensitive to the Orsay virus (3). In controlled infections, we commonly observed higher infection rates of *C. briggsae* by SANTV or LEBV than of *C. elegans* by ORV (e.g. respectively 60-80% versus 30-50% of the animals infected at 7 days after infection). The differences in infectivity between SANTV and LEBV on one hand and the ORV on the other hand could explain the lower probability to detect infected *C. elegans* than infected *C. briggsae*.

The *C. briggsae* species is distributed all over the world (16), yet we rarely found signs of infection outside France, and never outside Europe. Whether this geographical bias in viruses’ sampling reflects the geographical distribution of the sensitivity of *C. briggsae* to the noda-like viruses is still to be clarified. On another note, the time between the collection in the field and the processing of the sample (rotting vegetal matter) in the laboratory could impact the survival of the *Caenorhabditis* animals, especially for the infected animals, and therefore lower the probability to recover infected animals.

The Santeuil and Le Blanc viruses are often sympatric in France, co-existing even in the same individual worm and individual cell. They are thus in principle susceptible to exchange genetic material. Reassortment, by shuffling viral RNA molecules between various species, likely plays an important role in viral evolution (39, 40). The reassortment of RNA molecules (horizontal gene transfer) can occur between highly divergent viruses. For instance, the mosinovirus (MoNV) originated from the reassortment between a virus closely related to the Pariacoto virus (*Nodaviridae*) [MoNV RdRP shares 43% amino acid identity with Pariacoto virus] and a virus closely related to Lake Sinai virus 2 [MoNV capsid shares 16% amino acid with the LSV 2] (39). In our study, sequence analysis did not detect genetic exchange between the SANTV and LEBV, but did detect several reassortment events between SANTV variants.

*Caenorhabditis* nematodes and their noda-like viruses thus provide an exciting model to investigate co-evolution dynamics between an animal host and its natural viruses. First, *C. briggsae* and *C. elegans* are model organisms with a short life cycle, which eases multigenerational experiments and quantitative genetic analysis. Second, the natural combinations of one *Caenorhabditis* isolate with its natural viral variant can easily be isolated, maintained under laboratory conditions, kept frozen and revived. Third, natural virus-nematode combination can be dissociated by storing the viral isolate as a filtrate and by bleaching the *Caenorhabditis* isolate to remove viral infections (as in (21)). Fourth, the infection can be reconstituted by transgenesis (8), allowing to test the effect of precise sequence changes. Fifth, we developed here a phenotypic assay to test the competitive ability of two viruses in different host genotypes. This leads the way to further studies of phenotypically significant evolutionary change in both host and virus.

## Acknowledgements

We thank participants in field work, especially Sarah Marsh and Robert Luallen. M.-A.F. is grateful to the GENiE (Group of *Elegans* New Investigators in Europe) group for their invitation to Mělník. This work was funded by grants from the Agence Nationale de la Recherche (ANR-11-BSV3-013) and the Fondation pour la Recherche Médicale (DEQ20150331704) to MAF and supported in part by grants from National Institutes of Health (R01 AI134967 and R21AI119917) to DW.

## Supplemental data

**Figure S1. Tissue tropism of the Mělník virus in *C. briggsae*.** JU3272 and JU3276 *Cbr* isolates were naturally infected. The *Cbr* isolate JU1498 was artificially infected with MELV JUv3272. FISH staining of MELV-infected was performed using one 21-nt long probe (Texas Red) targeting the MELV RNA1 molecule. Nuclei were counterstained with DAPI (merge panels). h: Head of *C. briggsae* animals. Red: MELV RNA1 probe with 200ms exposition time; Grey: DAPI staining. Scale bars represent 100 μm.

**Figure S2. Amino-acid alignment of the RdRP protein of the four viruses.** The sequences of the new virus (MELV) are indicated in red.

**Figure S3. Amino-acid alignment of the caspid-delta protein of the four viruses.**

**Figure S4. Amino-acid alignment of the RdRP and Capsid-delta proteins of SANTV variants and MELV.** MELV sequences are in red, SANTV RNA1-clade A in black and SANTV RNA1-clade B in blue.

**Figure S5. Evolutionary relationships of MELV and SANTV variants. Left panel:** Phylogenetic relationships inferred using the Neighbor-Joining method (30) on the alignments of nucleotide sequences. The tree is drawn to scale with the number of base substitutions per site. **Right panel:** A timetree inferred using the Reltime method (29) and estimates of branch lengths inferred using the Neighbor-Joining method (30). Bars around each node represent 95% confidence intervals. Codon positions included were 1st+2nd+3rd+Noncoding. All positions containing gaps and missing data were eliminated. There was a total of 1709 positions in the final RNA1 dataset and 2251 positions in the final RNA2 dataset.

**Figure S6. Comparison of ORV variants’ virulence on different *C. elegans* wild isolates.** N2, JU1580, MY10 and JU2572 *C. elegans* (*Cel)* isolates were infected with the ORV variants JUv2572 and JUv1580. For each *C. elegans* isolate, five L4 larvae were infected with 50 µL of the ORV viral filtrate (either JUv1580 or JUv2572) and cultures were propagated for two/three generations at 20°C. Infections were performed in triplicates. ORV-infections were monitored on F2 adult hermaphrodites by FISH using Biosearch probe (Cal Fluor Red® 610) targeting the ORV RNA2 molecule. **A.** Proportion of infected F2/F3 adult hermaphrodites. The total number of animals scored are given below the histograms. Bars represent the standard deviation among replicates. ***, *: p<0.001, 0.05 in a glm taking the viral variant knowing the host genotype into account, each virus effect being compared to JUv1580 on the host *C. elegans* JU1580. **B.** Number of infected cells per infected F2/F3 adult hermaphrodite. The total number of infected animals are given below the histograms. **C/D.** Distribution of infected cells along *C. elegans* intestine (9 rings of intestinal cells, 1 being close to the pharynx and 9 to the rectum) considering all the *Cel* isolates or or each of them separately.

**Figure S7. Pairwise infections of *C. briggsae* JU1264 with one LEBV and two SANTV variants. (A)** Design of the assay for the pairwise competition between JUv1993 SANTV and either LEBV JUv1498 or SANTV JUv1551. The results of the above experiment in *C. briggsae* JU1264 are shown in panels B and C. Results are expressed as a fraction of infected individuals (top graphs) and mean number of infected cells in the infected animals (bottom graphs), across three generations. (Co-) infection levels were scored using FISH staining of infected cells/animals. **(B)** For the competition between SANTV variants we used single oligonucleotide probes targeting JUv1551 RNA1 (ATTO425) and JUv1993 RNA1 (Texas Red) (see Table S1c). **(C)** To score (co-) infection levels in the competition between SANTV and LEBV we used custom Stellaris™ (Biosearch Technologies) probes targeting JUv1264 RNA1 (Cal fluor Red 610) and JUv1498 RNA1 (Quasar 670) (see Table S1b).

**Table S1. List of oligonucleotides.**

a. List of oligonucleotides used for RT-PCR, PCR and Sanger sequencing
b. List of FISH probe sets
c. List of single oligonucleotide probes for FISH.

**Table S2. List of *Caenorhabditis* and viral strains.**

The colors correspond to the presence of the different viruses.

**Table S3. Probabilities of the reassortment events between viral variants detected using RDP4.** The *p*-value corresponds to the rejection of the hypothesis of absence of reassortment or recombination for the detected events.

## References

1. Félix M-A, Ashe A, Piffaretti J, Wu G, Nuez I, Bélicard T, Jiang Y, Zhao G, Franz CJ, Goldstein LD, Sanroman M, Miska EA, Wang D. 2011. Natural and Experimental Infection of Caenorhabditis Nematodes by Novel Viruses Related to Nodaviruses. PLoS Biol 9:e1000586.

2. Franz CJ, Zhao G, Felix M-A, Wang D. 2012. Complete Genome Sequence of Le Blanc Virus, a Third Caenorhabditis Nematode-Infecting Virus. J Virol 86:11940–11940.

3. Ashe A, Bélicard T, Le Pen J, Sarkies P, Frézal L, Lehrbach NJ, Félix M-A, Miska EA. 2013. A deletion polymorphism in the Caenorhabditis elegans RIG-I homolog disables viral RNA dicing and antiviral immunity. eLife 2.

4. Tanguy M, Véron L, Stempor P, Ahringer J, Sarkies P, Miska EA. 2017. An Alternative STAT Signaling Pathway Acts in Viral Immunity in *Caenorhabditis elegans*. mBio 8.

5. Jiang H, Chen K, Sandoval LE, Leung C, Wang D. 2017. An Evolutionarily Conserved Pathway Essential for Orsay Virus Infection of *Caenorhabditis elegans*. mBio 8.

6. Le Pen J, Jiang H, Di Domenico T, Kneuss E, Kosalka J, Leung C, Morgan M, Much C, Rudolph KLM, Enright AJ, O’Carroll D, Wang D, Miska EA. 2018. Terminal uridylyltransferases target RNA viruses as part of the innate immune system. Nat Struct Mol Biol 25:778–786.

7. Franz CJ, Renshaw H, Frezal L, Jiang Y, Félix M-A, Wang D. 2014. Orsay, Santeuil and Le Blanc viruses primarily infect intestinal cells in Caenorhabditis nematodes. Virology 448:255–264.

8. Jiang H, Franz CJ, Wu G, Renshaw H, Zhao G, Firth AE, Wang D. 2014. Orsay virus utilizes ribosomal frameshifting to express a novel protein that is incorporated into virions. Virology 450–451:213–221.

9. Fan Y, Guo YR, Yuan W, Zhou Y, Holt MV, Wang T, Demeler B, Young NL, Zhong W, Tao YJ. 2017. Structure of a pentameric virion-associated fiber with a potential role in Orsay virus entry to host cells. PLOS Pathog 13:e1006231.

10. Yuan W, Zhou Y, Fan Y, Tao YJ, Zhong W. 2018. Orsay δ Protein Is Required for Nonlytic Viral Egress. J Virol 92.

11. Bakowski MA, Desjardins CA, Smelkinson MG, Dunbar TA, Lopez-Moyado IF, Rifkin SA, Cuomo CA, Troemel ER. 2014. Ubiquitin-Mediated Response to Microsporidia and Virus Infection in C. elegans. PLoS Pathog 10:e1004200.

12. Chen K, Franz CJ, Jiang H, Jiang Y, Wang D. 2017. An evolutionarily conserved transcriptional response to viral infection in Caenorhabditis nematodes. BMC Genomics 18.

13. Reddy KC, Dror T, Sowa JN, Panek J, Chen K, Lim ES, Wang D, Troemel ER. 2017. An Intracellular Pathogen Response Pathway Promotes Proteostasis in C. elegans. Curr Biol 27:3544–3553.e5.

14. Kiontke KC, Félix M-A, Ailion M, Rockman MV, Braendle C, Pénigault J-B, Fitch DH. 2011. A phylogeny and molecular barcodes for Caenorhabditis, with numerous new species from rotting fruits. BMC Evol Biol 11:339.

15. Félix M-A, Duveau F. 2012. Population dynamics and habitat sharing of natural populations of Caenorhabditis elegans and C. briggsae. BMC Biol 10.

16. Félix M-A, Ailion M, Hsu J-C, Richaud A, Wang J. 2018. Pristionchus nematodes occur frequently in diverse rotting vegetal substrates and are not exclusively necromenic, while Panagrellus redivivoides is found specifically in rotting fruits. PLOS ONE 13:e0200851.

17. Barrière A, Felix M-A. 2014. Isolation of C. elegans and related nematodes. WormBook 1–19.

18. Richaud A, Zhang G, Lee D, Lee J, Félix M-A. 2018. The Local Coexistence Pattern of Selfing Genotypes in *Caenorhabditis elegans* Natural Metapopulations. Genetics 208:807–821.

19. Troemel ER, Félix M-A, Whiteman NK, Barrière A, Ausubel FM. 2008. Microsporidia Are Natural Intracellular Parasites of the Nematode Caenorhabditis elegans. PLoS Biol 6:e309.

20. Samuel BS, Rowedder H, Braendle C, Félix M-A, Ruvkun G. 2016. *Caenorhabditis elegans* responses to bacteria from its natural habitats. Proc Natl Acad Sci 113:E3941–E3949.

21. Stiernagle T. 2006. Maintenance of C. elegans. WormBook.

22. Zhao G, Wu G, Lim ES, Droit L, Krishnamurthy S, Barouch DH, Virgin HW, Wang D. 2017. VirusSeeker, a computational pipeline for virus discovery and virome composition analysis. Virology 503:21–30.

23. Schneider CA, Rasband WS, Eliceiri KW. 2012. NIH Image to ImageJ: 25 years of image analysis. Nat Methods 9:671–675.

24. Raj A, van den Bogaard P, Rifkin SA, van Oudenaarden A, Tyagi S. 2008. Imaging individual mRNA molecules using multiple singly labeled probes. Nat Methods 5:877–879.

25. Edgar RC. 2004. MUSCLE: multiple sequence alignment with high accuracy and high throughput. Nucleic Acids Res 32:1792–1797.

26. Kumar S, Stecher G, Tamura K. 2016. MEGA7: Molecular Evolutionary Genetics Analysis Version 7.0 for Bigger Datasets. Mol Biol Evol 33:1870–1874.

27. Librado P, Rozas J. 2009. DnaSP v5: a software for comprehensive analysis of DNA polymorphism data. Bioinformatics 25:1451–1452.

28. Jones DT, Taylor WR, Thornton JM. 1992. The rapid generation of mutation data matrices from protein sequences. Bioinformatics 8:275–282.

29. Tamura K, Battistuzzi FU, Billing-Ross P, Murillo O, Filipski A, Kumar S. 2012. Estimating divergence times in large molecular phylogenies. Proc Natl Acad Sci 109:19333–19338.

30. Saitou N, Nei M. 1987. The neighbor-joining method: a new method for reconstructing phylogenetic trees. Mol Biol Evol.

31. Martin DP, Murrell B, Golden M, Khoosal A, Muhire B. 2015. RDP4: Detection and analysis of recombination patterns in virus genomes. Virus Evol 1.

32. Nei M, Kumar S. 2000. Molecular Evolution and Phylogenetics. Oxford University Press. Oxford University Press Inc, New York, United States.

33. Tamura K, Nei M, Kumar S. 2004. Prospects for inferring very large phylogenies by using the neighbor-joining method. Proc Natl Acad Sci 101:11030–11035.

34. Pond SLK, Frost SDW, Muse SV. 2005. HyPhy: hypothesis testing using phylogenies. Bioinformatics 21:676–679.

35. Tamura K, Peterson D, Peterson N, Stecher G, Nei M, Kumar S. 2011. MEGA5: Molecular Evolutionary Genetics Analysis Using Maximum Likelihood, Evolutionary Distance, and Maximum Parsimony Methods. Mol Biol Evol 28:2731–2739.

36. Muse SV, Gaut BS. 1994. A likelihood approach for comparing synonymous and nonsynonymous nucleotide substitution rates, with application to the chloroplast genome. Mol Biol Evol.

37. Felsenstein J. 1981. Evolutionary trees from DNA sequences: A maximum likelihood approach. J Mol Evol 17:368–376.

38. Nei M, Gojobori T. 1986. Simple methods for estimating the numbers of synonymous and nonsynonymous nucleotide substitutions. Mol Biol Evol.

39. Schuster S, Zirkel F, Kurth A, van Cleef KWR, Drosten C, van Rij RP, Junglen S. 2014. A Unique Nodavirus with Novel Features: Mosinovirus Expresses Two Subgenomic RNAs, a Capsid Gene of Unknown Origin, and a Suppressor of the Antiviral RNA Interference Pathway. J Virol 88:13447–13459.

40. Lowen AC. 2018. It’s in the mix: Reassortment of segmented viral genomes. PLOS Pathog 14:e1007200.

